# Super Bloom: Fast and precise filter for streaming *k*-mer queries

**DOI:** 10.64898/2026.03.17.712354

**Authors:** Etienne Conchon-Kerjan, Timothe Rouzé, Lucas Robidou, Florian Ingels, Antoine Limasset

## Abstract

Approximate membership query structures are used throughout sequence bioinformatics, from read screening and metagenomic classification to assembly, indexing, and error correction. Among them, Bloom filters remain the default choice. They are not the most efficient structures in either time or memory, but they provide an effective compromise between compactness, speed, simplicity, and dynamic insertions, which explains their widespread adoption in practice. Their main drawback is poor cache locality, since each query typically requires several random memory accesses. Blocked Bloom filters alleviate this issue by restricting accesses for any given element to a single memory block, but this usually comes with a loss in accuracy at fixed memory.

In this work, we introduce the Super Bloom Filter, a Bloom filter variant designed for streaming *k*-mer queries on biological sequences. Super Bloom uses minimizers to group adjacent *k*-mers into super-*k*-mers and assigns all *k*-mers of a group to the same memory block, thereby amortizing random accesses over consecutive *k*-mer queries and improving cache efficiency. We further combine this layout with the findere scheme, which reduces false positives by requiring consistent evidence across overlapping subwords. We provide a theoretical analysis of the construction of Super Bloom filters, showing how minimizer density controls the expected reduction in memory transfers, and derive a practical parameterization strategy linking memory budget, block size, collision overhead, and the number of hash functions to robust false-positive control.

Across a broad range of memory budgets and numbers of hash functions, Super Bloom consistently outperforms existing Bloom filter implementations, with several-fold time improvements. As a practical validation, we integrated it into a Rust reimplementation of BioBloom Tools, a sequence screening tool that builds filters from reference genomes and classifies reads through *k*-mer membership queries for applications such as host removal and contamination filtering. This replacement yields substantially faster indexing and querying than both the original C++ implementation and Rust variants based on Bloom filters and blocked Bloom filters. The findere scheme also reduces false positives by several orders of magnitude, with some configurations yielding no observed false positives among 10^9^ random queried *k*-mers. Code is available at https://github.com/EtienneC-K/SuperBloom and https://github.com/Malfoy/SBB.

## 1. Introduction

Approximate membership data structures, often called filters, are widely used in large-scale database systems [1], [2], [3], storage systems [4], [5], [6], and networked services [7], [8], [9], [10]. Their appeal is simple: they can quickly rule out absent keys while using very little memory, which is especially valuable when the represented set contains a massive number of elements and the keys themselves are large.

Formally, a filter represents a set *S* ⊂ *U* of size *n* = |*S*|, where *U* is the universe of possible keys, and answers membership queries approximately. In exchange for compactness, filters allow a nonzero falsepositive rate *ε*. Most practical filters support constant-time queries, and in the usual sparse regime, where |*U* | ≫ *n*, any approximate membership data structure requires at least 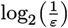 bits per key [11].

The most widely used filter is the Bloom filter. Bloom filters are simple to implement and can achieve space usage within a factor 1.44 of the information theoretic optimum, which corresponds to 44% overhead over the lower bound [7]. They also support dynamic insertions, which makes them a robust default choice across many applications.

Because sequence alignment is computationally expensive, many bioinformatics methods switched to alignment-free heuristics [12], [13], [14]. These methods typically compare sequences through their sets of *k*-mers, that is, substrings of fixed length *k*, and use *k*-mer similarity as a proxy for sequence relatedness [15], [16], [17]. This makes it possible to process sequences in linear time in settings where exact alignment would be too costly. Their main drawback, however, is memory usage, since representing a sequence as a set of *k*-mers increases the memory footprint by roughly a factor of *k* compared with storing the raw sequence alone. To reduce this cost, sequence bioinformatics makes extensive use of memory-efficient probabilistic data structures such as Bloom filters for storing *k*-mers. This is usually acceptable because a low false-positive rate has a limited impact on *k*-mer-based methods: unrelated sequences may share a few spurious hits without being considered genuinely similar[18].

As a result, Bloom filters are used throughout the bioinformatics software ecosystem, including:

- Large-scale sequence and dataset indexing based on *k*-mers or minimizers, such as BIGSI [19], COBS [20], PAC [21], HowDeSBT [22], SBT [18], SSBT [23], AllSomeSBT [24], Raptor [25], RAMBO [26], MetaProFi [27], kmtricks [28], BFT [29], Needle [30], and HIBF [31]
- Accelerating read mapping, such as DREAM-Yara [32]
- Genome assembly and gap filling, such as ABySS [33], Minia [34], RNABloom [35], RNA-Bloom2 [36], and Sealer [37]
- Metagenomic classification, such as ganon [38], ganon2 [39], and KMCP [40]
- *k*-mer counting, such as khmer [41], BFCounter [42], and Turtle [43]
- Error correction, such as BLESS[44], BLESS2[45], Bloocoo [46], BFC [47] Lighter [48], LoRDEC [49], and Lorma [50]
- Paired-end connection, such as Konnector [51] and ABySS-Connector [52]
- Read compression, such as Leon [53] and BARCODE [54]
- Transcript quantification, such as RNA-Skim8 [55] and Reindeer2 [56]
- Variant and splicing detection, such as DiscoSNP [57], DiscoSNP+ + [58], MindTheGap [59], FusionBloom [60], and KisSplice [61]
- Genome polishing, such as ntEdit [62]
- Sequence selection, such as aKmerBroom [63] and BioBloom Tools [64]
- Differential analysis, such as kmdiff [65]
- Compacted de Bruijn graph construction, such as TwoPaCo [66] and Bifrost [67].

Despite this ubiquity, their use is often suboptimal for several reasons. A first limitation is that a Bloom filter requires multiple hash functions to achieve a low false-positive rate. With *m* bits storing *n* keys, the false positive rate is minimized for 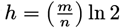 hash functions, which is approximately 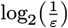 when tuned for a target false positive rate *ε* [68]. When the filter is large, this translates into multiple costly random memory accesses per query. A common practical workaround is to cap the number of hash functions to improve speed, but this increases the space required to achieve a given false positive rate. Blocked Bloom filters[69] address this issue by constraining all hashes for a key to a small memory block, ensuring that each query performs a single random access regardless of the number of hashes. This restores time efficiency at a moderate space cost, around 56% overhead instead of about 44% for optimal Bloom filters, with only a small increase in implementation complexity. Despite this attractive tradeoff, blocked variants remain underused in most bioinformatics pipelines.

A second limitation is that Bloom filters pay a space cost for being dynamic. Static filters, built once for a fixed set of keys and queried afterwards, can be substantially more space efficient. In particular, practical static filters such as Fuse filters [70], which improve over XOR-based designs[71], [72], can reach overheads as low as about 8.5% while providing fast queries with a single random access and very fast construction, below 100ns per key. Yet, few bioinformatics tools benefit from these structures, with notable exceptions such as Taxor [73]. This gap matters because many bioinformatics workflows naturally separate construction and querying into two phases, with no interleaving of insertions and queries, so a static filter would bring only benefits. While reducing overhead from 56% to 8.5% may appear modest, its impact increases with the bits per key budget. For 4 bits per key, this corresponds to about a 2.19× reduction in false positive rate, for 8 bits per key about 4.79×, and for 16 bits per key about 23×. More recent designs, such as bumped ribbon filters [74], built on ribbon filters [75], achieve near-optimal space usage, with overhead below 1% and in some configurations as low as 0.1%.

A third limitation is that standard Bloom filters are not adaptable. If more elements than expected are inserted, the false positive rate increases until the filter becomes ineffective and answers yes to almost everything. In practice, the filter must be sized in advance based on an expected cardinality, so datasets that vary widely in size end up with markedly different false positive rates under the same parameterization. More broadly, Bloom filters do not provide strong guarantees when the inserted set size deviates from the design target. This has motivated a large body of work on dynamic and scalable alternatives. Scalable Bloom filters[76] grow by adding new subfilters while enforcing a global false positive target, quotient filters [77] improve locality and support deletions via compact quotienting, Morton filters [78] are faster than Cuckoo Filter [79] through biasing and compression, vacuum filters [80] reduce space and improve throughput using semi-sorting and locality, InfiniFilter [81] expands by growing the table and lengthening fingerprints over time, and Aleph filters [82] target constant-time operations under unbounded growth while retaining static-like tradeoffs when a growth estimate is available.

However, depending on the usage, these improvements are not always directly applicable. Some usage requires a dynamic filter, some can have reasonable cardinality estimates using sequence length proxies ignoring repeated *k*-mers, and some use inverted layouts where many per-dataset filters are stored transposed[20], [21]. In that setting, a *k*-mer query reads the *h* bit-slices corresponding to its hash positions and intersects them to obtain candidate datasets, so the query time is driven by slice bandwidth and bitwise operations rather than by per-filter random accesses. Determining which filter designs best match these bioinformatics access patterns remains an open optimization problem.

Recently, several works have noted that classical lower bounds can be improved in bioinformatics by exploiting the fact that *k*-mers are not independent [83], [84], [85]. One natural way to capture this dependence is through minimizers. Given a *k*-mer, a minimizer is an *m* -mer selected from among its constituent substrings of length *m*, and the rule used to choose that *m*-mer is called the minimizer scheme. Although many minimizer schemes[86], [87], [88], [89], [90] have been proposed, the simplest and most widely used approach is to order candidate *m*-mers by hash value and, for each *k*-mer, to select the one with minimum hash value. In bioinformatics applications, sequences are typically parsed sequentially with a sliding window of length *k*, so consecutive *k*-mers overlap by *k* − 1 symbols. As a result, many adjacent *k*-mers share the same minimizer and can therefore be grouped into super-*k*-mers, making explicit the strong local structure of sequence-derived *k*-mer sets. This structure is particularly useful for building minimal perfect hash functions, which assign each distinct *k*-mer a unique identifier and constitute a standard primitive for compact exact dictionaries and for associating satellite information to *k*-mers[91], [92], [93], [94], [95]. LPHash [85] exploits this minimizer-based organization by first hashing distinct minimizers, then recovering the identifiers of the *k*-mers inside each super-*k*-mer from their relative positions with respect to the shared minimizer. More precisely, once the minimizer of a query *k*-mer is known, LPHash uses the identifier of that minimizer to locate the corresponding super-*k*-mer and derives the rank of the query *k*-mer from its local position within the block. In this way, consecutive *k*-mers tend to receive consecutive identifiers, yielding a locality-preserving MPHF rather than an arbitrary one. LPHash further reduces the space cost by encoding super-*k* -mers according to the position of the minimizer within their boundary *k*-mers, so that part of the offset information becomes implicit, while the relatively rare ambiguous minimizers that occur in multiple super-*k*-mers are handled separately. Altogether, this makes it possible to go below the usual 1.44 bits-per-key bound for independent inputs, with practical space usage below 1 bit per *k*-mer. LPHash illustrates how overlap can be exploited structurally to improve dictionaries. A complementary line of work exploits the same dependence at query time in approximate membership structures, where a *k*-mer is validated through the correlated evidence carried by its overlapping subwords. The meta-technique findere [83] indexes *s*-mers with *s* < *k* in the underlying filter and answers *k*-mer queries by scanning overlapping *s* -mers along the query sequence. It reports a *k*-mer hit only when the implied *s*-mer answers contain a stretch of at least *z* with *z* = *k* − *s* + 1 consecutive positives. Since false positives are unlikely to form long consecutive runs, the effective false positive rate drops roughly like *ε*^*z*^ for a base *s*-mer false positive rate *ε*. This false positive rate reduction comes at the cost of adding *z* − 1 insertions (resp. query) in the filter per indexed (res. queried) read.

In this paper, we propose an efficient filtering structure that exploits the streaming nature of sequence-derived insertions and queries and make three contributions.

First, we introduce the ***SBF*** (Super Bloom Filter), a Bloom filter tailored for streaming *k*-mer queries. Inspired by blocked Bloom filters, SBF exploits the structure of super-*k*-mers by assigning all *k*-mers that share minimizers to the same memory block. This changes the access pattern from one random memory access per *k*-mer to one random access per super-*k*-mer, improving cache locality and reducing memory-bandwidth pressure. As the expected length of a super-*k*-mer grows with *k* for fixed *m*, this locality benefit already appears for practical values such as *k* = 31 and becomes stronger for larger *k* values.

Second, we adapt the findere technique to this super-*k*-mer setting. This adaptation substantially reduces false positives while preserving a tunable sensitivity to alien *k*-mers that remain similar to indexed *k*-mers. We further analyze the associated memory trade-offs and show that optimal memory usage can be achieved for any choice of the findere parameter. This makes it possible to control the balance between false-positive filtering and tolerance to near matches without sacrificing the simplicity of the underlying Bloom filter design.

Third, we provide an efficient implementation of our approach (https://github.com/EtienneC-K/SuperBloom) and evaluate it against other high-performance Bloom filter implementations. As a practical demonstration, we also integrate our method into a Rust fork of BioBloom Tools[64], a Bloom-filter based sequence screening tool that builds filters from reference genomes and uses *k*-mer membership queries to rapidly classify reads for applications such as host removal and contamination filtering. Replacing its standard Bloom filters with our variant yields improved throughput together with better falsepositive control, showing that the method is not only theoretically well motivated but also directly useful in real bioinformatics workflows. This reimplementation is available at https://github.com/Malfoy/SBB.

## 2. Methods

We consider the indexing of a set of *k*-mers using three related structures, namely the Bloom filter, the blocked Bloom filter, and the proposed Super Bloom Filter (***SBF***). Throughout the section, *n* denotes the number of indexed *k*-mers, *M* the total number of bits in the filter, and *k* the *k*-mer length. This theoretical analysis focuses on insertion and query costs, with particular attention to the number of memory transfers. Hash-computation costs are neglected in the cost model because they are negligible relative to random accesses in practice.

### 2.1. Bloom filters

A Bloom filter stores a set of keys in a bit-vector of length *M* using *h* hash functions *H*_1_, …, *H*_*h*_. Initially, all bits are set to 0. Inserting a *k*-mer *x* sets the *h* bits addressed by

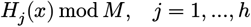

to 1. Conversely, a query returns that a *k*-mer is present if and only if all corresponding bits are equal to 1.

When querying an alien *k*-mer, it is possible that its *h* corresponding bits were set to 1 by a combination of inserted keys, resulting in a false positive. Under the standard independence approximation, after inserting *n k*-mers, the false-positive rate is

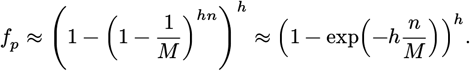

The number of hash functions that minimize *f*_*p*_ is 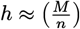 ln 2, which Yields 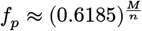.

From a computational perspective, each insertion or query performs *h* accesses to unrelated positions in the bit-vector. Neglecting hash-computation costs, the per-*k*-mer cost can be summarized as

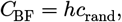

where *c*_rand_ is the cost of one random memory access. In practice, the main bottleneck of the classical Bloom filter is this random memory-access pattern, as each operation performs *h* scattered accesses with poor spatial locality.

### 2.2. Blocked Bloom filters

A blocked Bloom filter partitions the bit-vector into *B* blocks of *b* bits, with *M* = *B* · *b*. A first mapping chooses a block, and the *h* Bloom filter hashes select bit positions only inside that block. For a *k*-mer *x*, let *g*(*x*) ∈ {0, …, *B* − 1} denote the block index and let

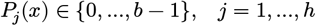

denote the *h* local positions inside the selected block. Insertion sets to 1 the bits (*g*(*x*), *P*_1_(*x*)), …, (*g*(*x*), *P*_*h*_(*x*)) and a query checks the same positions.

If the blocks are selected uniformly, each block receives on average

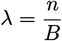

*k*-mers. Under the same approximation as above, the false-positive rate of a typical block is

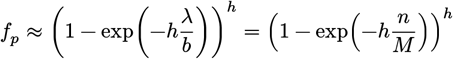

At first order, this matches the classical Bloom filter while changing the memory-access pattern.

The main advantage of the block organization is that all the bits accessed for one *k*-mer insertion/query are confined to a single block. If one block fits in one cache line or in a small fixed number of machine words, the whole insertion or query requires only one memory access. Neglecting hash-computation costs, the per-*k*-mer cost becomes

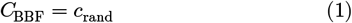

Compared with the classical Bloom filter, the number of random memory transfers drops from *h* scattered accesses to a single block access per *k*-mer.

### 2.3. Minimizers and super-*k*-mers

Our proposal keeps the block-based organization of blocked Bloom filters, but changes the object used to dispatch data to blocks. Instead of assigning each *k*-mer independently to a block, we group consecutive *k*-mers sharing the same minimizer into super-*k*-mers. Then, we assign all *k*-mers of a super-*k*-mer to the same block.

Let *m* < *k* be the minimizer length and let *w* = *k* − *m* + 1 be the number of overlapping *m*-mers contained in one *k*-mer. Given a random total order on all *m*-mers, the minimizer of a *k*-mer *x*, denoted by min(*x*), is the smallest of its *w* constituent *m*-mers under that order.

Along a sequence, consecutive *k*-mers often share the same minimizer. A *super-k-mer* is defined as a maximal run of consecutive *k*-mers having the same minimizer. If a sequence contains *n k*-mers and these *k*-mers are partitioned into *S* super-*k*-mers, then the particular minimizer density (i.e. the fraction of selected minimizers among all *m*-mers) is defined as

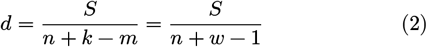

For random minimizers on random sequences, the expected density is well approximated [96] by

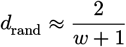

Equivalently, the expected number of *k*-mers per super-*k*-mer is 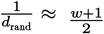 [97, Lemma 2]. This implies that minimizers induce runs of consecutive *k*-mers that can be processed together.

### 2.4. Super Bloom filters

The Super Bloom Filter keeps the local update rule of a blocked Bloom filter but changes the granularity of block assignment. A blocked Bloom filter assigns each *k*-mer independently to a block. In SBF, an entire super-*k*-mer *σ* = (*k*_1_, …, *k*_*r*_) is assigned to a single block through its shared minimizer min(*σ*). A mapping *g*(min(*σ*)) ∈ {0, …, *B* − 1} selects that block, and each constituent *k*-mer *k*_*i*_ is then inserted within the block using the same *h* local hash functions.

The local bit-setting rule is therefore unchanged. The difference is that the cost of loading the target block is paid once per super-*k*-mer instead of once per *k*-mer, which amortizes the random-access cost across consecutive sequence-derived queries.

For a stream of *n* consecutive *k*-mers partitioned into *S* super-*k*-mers, and neglecting hash-computation costs, the total insertion cost is

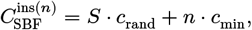

where *c*_rand_ is the cost of loading one target block and *c*_min_ is the amortized cost of computing minimizers along the sequence. The key difference from a blocked Bloom filter is that the block-loading cost is paid once per super-*k*-mer rather than once per *k*-mer. The same decomposition holds for queries,

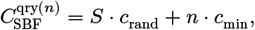

with local updates replaced by local bit tests.

Dividing by *n*, together with Equation 2, gives the amortized per-*k*-mer cost

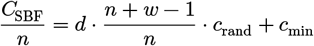

Using the random-minimizer approximation, and assuming *n* is large enough so that 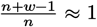,

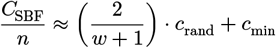

This expression makes the gain explicit. Relative to a blocked Bloom filter, recall Equation 1, the dominant block-transfer term is no longer paid once per *k*-mer but once per super-*k*-mer. Since the expected number of *k*-mers per super-*k*-mer is about 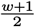, the expected number of block transfers per *k*-mer is reduced from 1 to 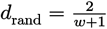; or, equivalently, the expected reduction factor on block loads is 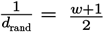.

In summary, the three structures exhibit the following asymptotic behavior per indexed *k*-mer:

- The Bloom filter requires *h* scattered accesses.
- The blocked Bloom filter reduces this to one block access.
- The Super Bloom filter further reduces the effective block-access rate to the minimizer density *d*, which is about 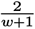 under the random-minimizer model.

Therefore, the proposed structure preserves the cache-local organization of blocked Bloom filters while exploiting super-*k*-mer coherence to amortize block loading over multiple consecutive *k*-mers.

It is important to note that this amortization applies to sequence-wise insertion and sequence-wise querying, which are the relevant regimes for *k*-mer indexing and querying in genomic data. For isolated *k*-mer queries taken independently, the Super Bloom filter still requires one minimizer computation and one block access per queried *k*-mer, so the cost then reverts to the same order as a blocked Bloom filter.

### False-positive rate and parameter selection

The block-wise false-positive rate can be made explicit by conditioning on the size of the super-*k*-mer assigned to a block. Consider a block of size *b* bits that receives a single super-*k*-mer *σ* = (*k*_1_, …, *kr*) containing *r k*-mers. Since each *k*-mer contributes *h* local bit updates inside that block, the total number of in-block insertions is *h* · *r*.

Under the standard approximation, the probability that a given bit remains equal to 0 after inserting *σ* is

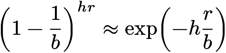

Therefore, the expected fraction of bits set to 1 in the block is

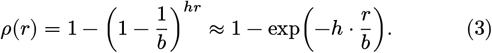

A negative *k*-mer mapped to that block yields a false positive when its *h* tested positions all fall on bits already set to 1. Conditionally on the super-*k*-mer length *r*, the block-wise false-positive rate is therefore

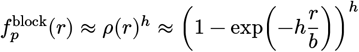

Note that this expression increases with *r*. Hence, large super-*k*-mers are the critical case, because they fill their target block more densely and therefore dominate the false-positive behavior. More generally, if a block receives several super-*k*-mers with sizes *r*_1_, …, *rt*, the same approximation holds by using *r* = *r*_1_ + … + *rt*. Thus the relevant worst-case quantity is not the average super-*k*-mer size, but the largest total number of indexed *k*-mers that may accumulate in one block.

For this reason, choosing *h* from the expected super-*k*-mer length is not robust. A super-*k*-mer that is significantly larger than average can overfill its block and sharply increase the false-positive rate. Instead, we are interested in the maximal number of indexed *k*-mers that may be inserted into one block, which we denote by *r*_max_. The maximum number of *k*-mers per super-*k*-mer is *w* (neglecting rare duplicate minimizer cases) ; therefore if we denote by *t*_max_ the maximum number of super-*k*-mers per block, we have

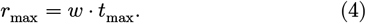

A robust choice of the number of hash functions is therefore obtained by imposing the half-full condition [98] on the worst admissible block

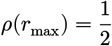

Together with Equation 3, we obtain

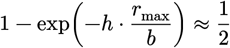

hence *h*_robust_ · *t*_max_ · *w* ≈ *b* ln 2. Equivalently,

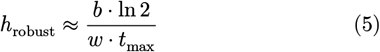

With this choice, the worst admissible block is half full on average. For every lighter block load *r* ≤ *r*_max_, one then has 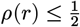 Therefore, for every admissible block load,

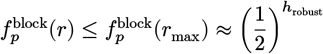

This choice is conservative rather than average-case optimal. It guarantees that even the heaviest admissible block remains at the standard Bloom-filter operating point. Smaller super-*k*-mers then operate below half filling and therefore have a lower false-positive rate than the worst-case design target.

The block size can be derived from the total memory budget and the expected number of super-*k*-mers. Assume the number of indexed *k*-mers is *n*. Under the random-minimizer model, the expected number of super-*k*-mers is evaluated [97, Lemma 1] as

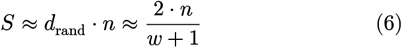

To control collisions between super-*k*-mers and blocks, we introduce an overhead parameter

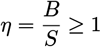

where *B* is the number of blocks. This parameter reflects the fact that for sufficiently large minimizer length *m*, repeated minimizers are rare for the overwhelming majority of buckets, as argued in SSHash [95], so the dominant residual source of block overloading is the collision of distinct super-*k*-mers into the same block. Equivalently, the expected number of super-*k*-mers per block is

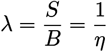

Assuming that super-*k*-mers are assigned independently and uniformly to blocks, the number of super-*k*-mers received by a block is well approximated by a Poisson law of parameter λ [69]. Therefore, the expected number of empty blocks is

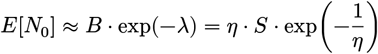

whereas the expected number of blocks receiving multiple super-*k*-mers, that is, at least two, is

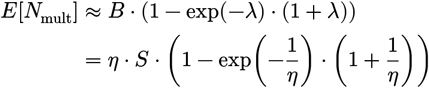

We tested our Poisson approximation in the Appendix, Section A.

Thus, increasing *η* reduces the fraction of colliding blocks, at the cost of creating more empty blocks.

Since the total filter size is *M* = *B* · *b*, with Equation 6, the corresponding block size is

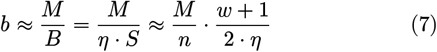

Substituting this value of *b* into Equation 5 yields

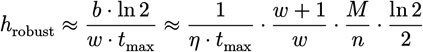

Thus, once *M, w*, and the input size *n* are fixed, the remaining design parameters are the overhead factor *η* and the maximal admissible number of super-*k*-mers per block *t*_max_. Increasing *η* decreases the collision rate but also decreases the block size. Increasing *t*_max_ forces a smaller number of hash functions, because the worst block must remain below half filling.

These expressions provide a practical parameterization rule for the Super Bloom filter. Given *M, k, m* and the input size *n*, with *w* = *k* − *m* + 1, one first estimates the expected number of super-*k*-mers from the minimizer density. One then chooses an overhead factor *η* to control the expected number of empty and colliding blocks, derives the block size from the memory budget, and finally sets *h* from the worst admissible number of super-*k*-mers per block *t*_{max}_.

However, when the memory budget becomes large, the corresponding block size *b* also grows, and the optimal number of hash functions increases accordingly. Past some point, additional memory therefore brings diminishing returns in practice: larger blocks are less likely to fit in cache, and the cost of computing and probing many hash functions is no longer negligible. Since the main motivation of the design is to preserve fast queries and updates, it is often preferable in practice to cap the block size at a reasonable value, for instance to the size of a cache line, even if the theoretical optimum would suggest a larger block. Although such a cap may appear suboptimal from a purely memory-theoretic perspective, it is not entirely detrimental, since using more blocks also reduces the probability that multiple super-*k*-mers are assigned to the same block.

### 2.6. Findere scheme implementation

To further reduce the false-positive rate, we adapt the findere principle at the block level. Instead of inserting the *k*-mers of a super-*k*-mer into its target block, we insert all of its constituent *s*-mers, with *s* < *k*. At query time, a *k*-mer is declared present in the block if and only if all of its constituent *s*-mers are present in that block.

Let *z* = *k* − *s* so that each *k*-mer contains *z* + 1 = *k* − *s* + 1 overlapping *s*-mers. Consider a super-*k*-mer *σ* = (*k*_1_, …, *k*_*r*_) containing *r k* -mers, assigned to a block of size *b* bits. The sequence spanned by *σ* has length *k* + *r* − 1, hence the number of overlapping *s*-mers inserted into the block is

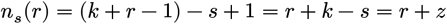

Assuming these *s*-mers are distinct and using the standard independence approximation, the probability that a given bit remains equal to 0 after insertion is

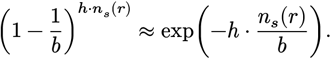

Therefore, the expected fraction of bits set to 1 in the block is

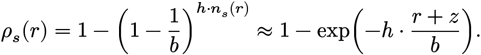

A queried *s*-mer that was not inserted in the block is reported positive with probability

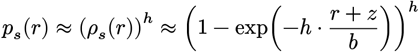

A random negative *k*-mer associated with that block contains *k* − *s* + 1 = *z* + 1 queried *s*-mers. Under the independence approximation, its false-positive probability is therefore

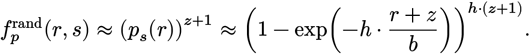

Thus, compared with direct *k*-mer insertion, the false-positive probability is exponentially reduced because all *z* + 1 constituent *s*-mers must simultaneously be positive.

There is, however, an analogue of the construction of false positives described for findere. Indeed, a negative *k*-mer may still share some of its *s*-mers with truly indexed sequence content, which increases its probability of being reported positive. An important limiting case is an *almost indexed k*-mer that differs from an indexed *k*-mer only by its last nucleotide. Such a *k*-mer shares its first *z s*-mers with the indexed *k*-mer, and only its last *s*-mer is different. Hence, its false-positive probability is simply

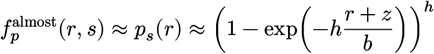

This is much larger than 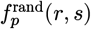, since only one constituent *s*-mer needs to be falsely recognized. We go over this point in greater detail empirically in Appendix Section B

As in the previous section, choosing *h* from the expected super-*k*-mer length is not robust. A long super-*k*-mer inserts many more *s*-mers than an average one and can therefore push its block far beyond the nominal operating point. Since each super-*k*-mer contributes at most to *w* + *z s*-mers in a block, the maximum number of inserted *s*-mers per block is therefore expressed as

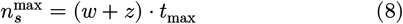

where *t*_max_, as in Equation 4, denotes the maximum number of super-*k*-mers allowed per block.

To compute a robust choice of the number of hash functions in such a case, we replace *r*_max_ by 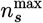 in Equation 5, which allows us to obtain

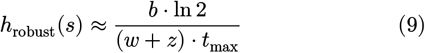

Using this robust choice, we obtain the following bound 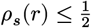 which allows us to derive

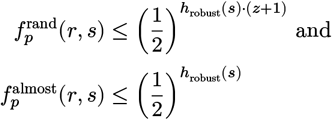

Using the same block size as previously, i.e. (cf. Equation 7)

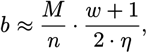

and substituting into Equation 9 yields

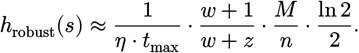

This analysis shows that, once *b* is dimensioned from the worst admissible block load, the parameter *s* mainly controls a trade-off between two kinds of false positives. When *s* decreases, a random negative *k* -mer must pass more constituent *s*-mer tests, so its false-positive probability decreases. At the same time, *h* decreases, so almost indexed *k*-mers that share all but one *s*-mers with truly indexed sequence content see their false-positive probability increasing. Therefore, *s* must be tuned according to the application. If preserving the distinction between related *k*-mers is desirable, a larger *s* may be beneficial.

It is also worth noting that, in the original findere setting[83], short *s* -mers may occur by chance elsewhere in the indexed data. As a consequence, taking *s* too small eventually leads to diminishing returns, because the stronger filtration obtained from requiring more constituent *s*-mers is progressively compensated by the increased probability of incidental *s*-mer matches. In the present setting, however, each block indexes only a very short sequence segment rather than a large collection of unrelated sequences. Accordingly, chance matches of short *s* -mers should remain much less frequent, and this effect is expected to remain negligible as long as 4^*s*^ ≫ *w*.

### 2.7. SuperBioBloom Tools

BioBloom Tools [99] is a Bloom-filter-based sequence screening framework designed to build filters from reference sequences and then classify sequencing reads according to filter membership. Its intended use is not full alignment but fast pre-filtering and categorization, especially for tasks such as host screening, contamination detection, and other quality-control steps where mapping positions are unnecessary. In that regime, Bloom-filter queries provide a substantially cheaper alternative to alignment while maintaining classification accuracy.

This is a natural application for our Rust reimplementation, since BioBloom Tools can directly benefit from alternative Bloom-filter layouts, namely blocked Bloom filters, and the proposed Super Bloom filters. It is also a setting in which the dynamic enrichment of the filter is well justified. Indeed, once a subset of reads or sequences has been selected as relevant to the target, their *k*-mers can be inserted into the filter as well, to broaden the representation of the target set and improve subsequent recall. This is particularly appropriate in screening applications, where the goal is not an exact set indexing but a sensitive retrieval of sequences related to the target.

### 2.8. Implementation details

Our implementation is designed for high-throughput sequence streaming. FASTQ/A parsing relies on needletail while minimizer computation uses SIMDMinimizers[100]. Consecutive *k*-mers and *s*-mers are hashed with rolling hashes based on ntHash[101], which avoids recomputing hash values from scratch at each step of the sequence scan. Membership queries are also short-circuited whenever possible. In the standard Bloom setting, probing stops as soon as one required bit is absent. In the findere variant, a queried *k*-mer is rejected as soon as one of its constituent *s*-mers is not found. This early stopping avoids unnecessary memory accesses and hash evaluations on negative *k*-mers, which are frequent in screening workloads. At the sequence level, SuperBioBloom Tools also benefits from an analogous short-circuiting strategy: when classifying a query sequence against a threshold on matching *k*-mers, the scan can stop as soon as the threshold is reached, but also as soon as it becomes impossible for the remaining unprocessed *k*-mers to reach it. Together, these early termination mechanisms further reduce the practical query cost, especially for clearly positive or clearly negative sequences.

When the expected cardinality is not provided, the implementation can perform a preliminary HyperLogLog-based estimation pass in order to dimension the filter automatically. This estimator is designed to remain lightweight relative to the full construction pipeline and is highly inspired by the SimdSketch Library (SimdSketch).

To reduce arithmetic overhead, the current implementation uses power-of-two sizes for the global filter and for the number of blocks. In that setting, block selection and local addressing can be implemented with inexpensive bit-mask operations instead of general modulo computations. Supporting arbitrary sizes would be possible using fast modulo techniques [102] and is left for future work.

Parallel construction relies on two levels of parallelism. Input data are first distributed across FASTQ/A chunks, and each chunk is then subdivided into sequence subchunks in order to improve load balancing across heterogeneous sequence lengths. To reduce allocation overhead, each worker thread reuses preallocated scratch buffers for temporary address vectors and related transient state. Synchronization during insertions is sharded over independent Bloom partitions, each protected separately, which substantially reduces lock contention compared with a single shared structure.

As mentioned in Section 2.4, our implementation divides the memory into blocks in which all the *k*-mers of a super-*k*-mer will be indexed. The indexing of a super-*k*-mer in a block, without using the findere scheme, is shown in Figure 1. Figure 2 shows the indexing of the same super-*k*-mer using the findere scheme. Figure 3 shows the query of a single present *k*-mer in the filter. Figure 4 shows the query of an alien *k*-mer. Under the assumption that it does not share any *s*-mer with an indexed *k*-mer, it would require 3 false-positive *s*-mers to yield a false positive for this *k*-mer. However, that assumption is not true for every alien *k*-mer. Figure 5 shows the query of a *k*-mer sharing all but one of its *s*-mers with an indexed *k*-mer. In this case, the false positive rate of such a *k*-mer is about the same as without the findere scheme (neglecting the small overhead introduced by the indexing of *k* − *s* more elements when using the findere scheme).

**Figure 1.**
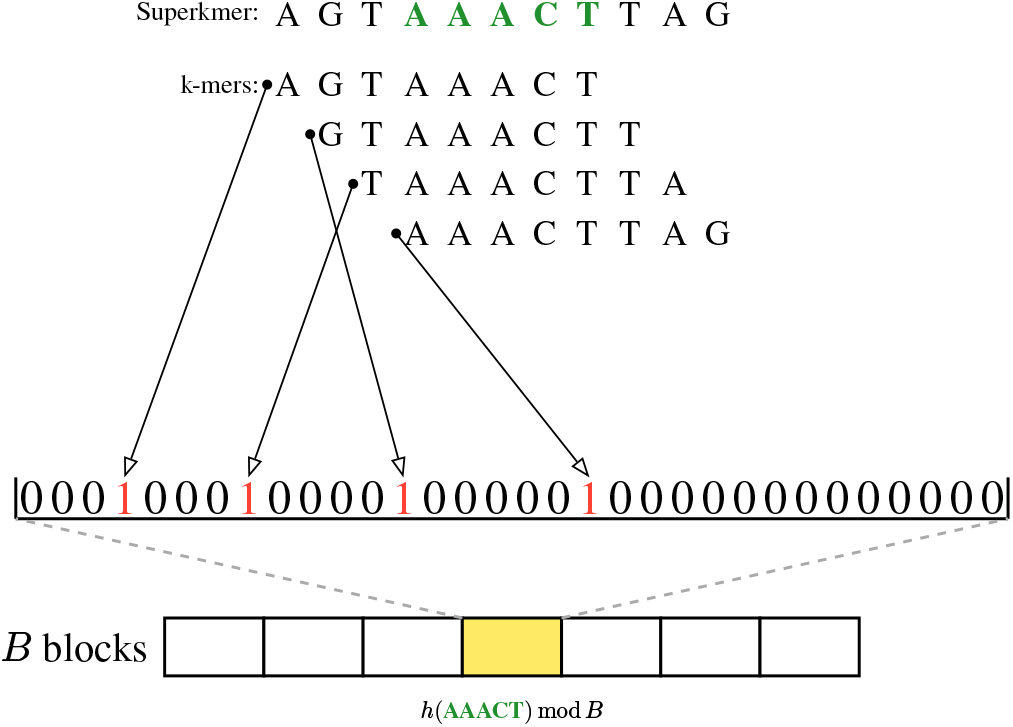
Indexing of a super-*k*-mer into a block (*k* = 8, *m* = 5, *B* = 7). The hash of its minimizer is used to select a block, in which its *k*-mers are inserted, guaranteeing they are all mapped to positions close in the memory.

**Figure 2.**
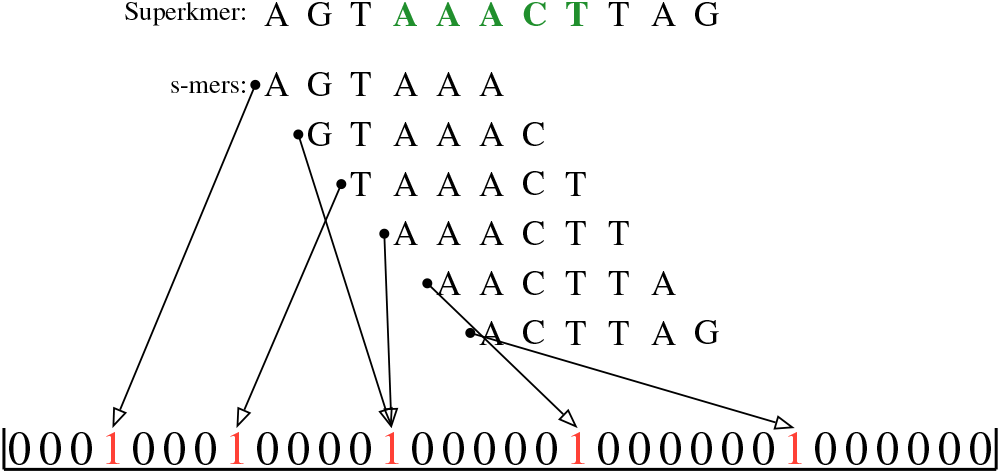
Indexing using the findere meta-scheme (*k* = 8, *m* = 5, *B* = 7, *s* = 6). During insertion, the *s*-mers of the super-*k*-mer are inserted into the filter.

**Figure 3.**
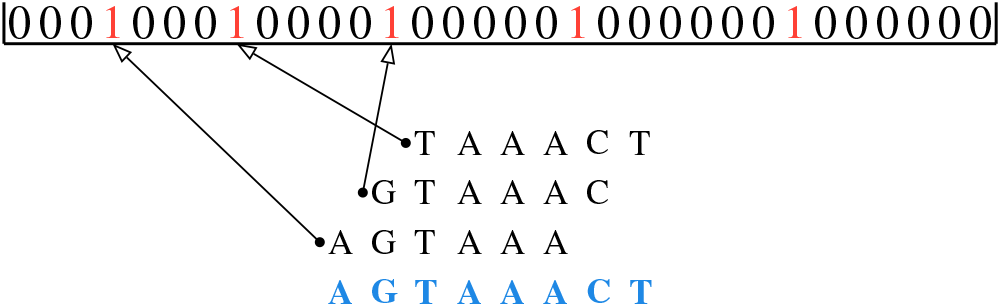
Querying an indexed *k*-mer with findere. The *s*-mers of the queried *k*-mer are searched in the filter. A *k*-mer is said to be present if and only if all its *s*-mers are found in the filter. Here, since all its *s*-mers are found in the filter, the *k*-mer is reported present.

**Figure 4.**
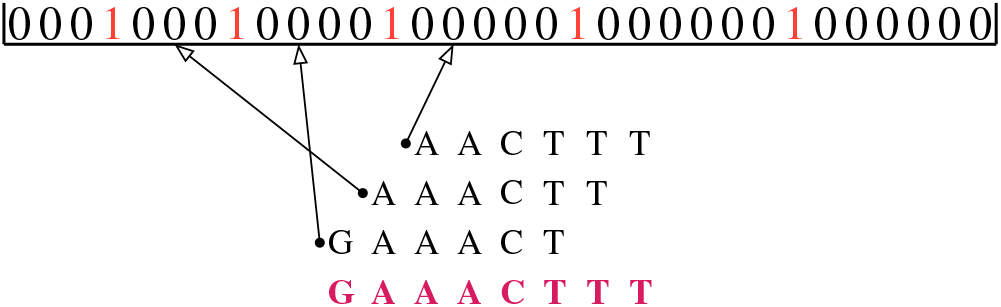
Querying an alien *k*-mer with findere. Its *s*-mers are not indexed, and it is likely that at least one of them will not be found, allowing the filter to reject the alien *k*-mer.

**Figure 5.**
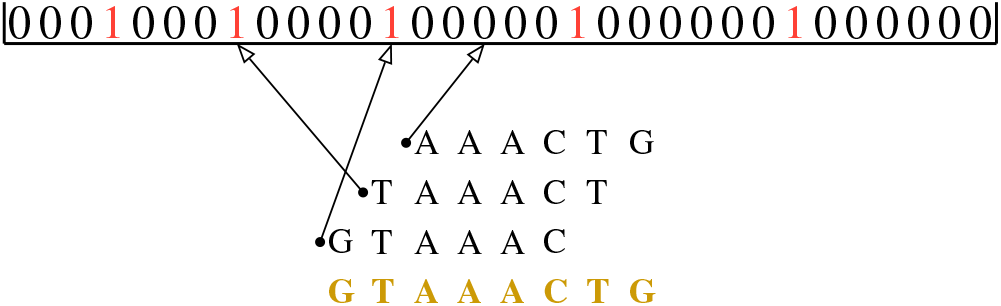
Querying with findere an alien *k*-mer sharing some *s*-mers with an indexed *k*-mer (namely “GTAAAC” and “TAAACT”). These *s*-mers will be found, as they were inserted. Thus, compared to a random alien *k*-mer, the filter will rely on fewer absent *s*-mers to reject it, increasing the probability of wrongly finding it.

## 3. Results

### 3.1. Application to BioBloom Tools

For all the following experiments, if not specified otherwise, *k* = 31, *m* = 21. Super Bloom was consistently the fastest layout in the BioBloom Tools setting. Using the human reference genome for indexing and 10× simulated human 10 kb reads for querying, CPU time increased with the number of hash functions for all methods, but much more slowly for blocked layouts and slowest for Super Bloom (Fig. Figure 6). At *h* = 10, Super Bloom indexed in less than 10^3^ s, compared with about 1.2×10^3^ s for the blocked Bloom filter, about 2.0×10^3^ s for the original C++ implementation, and about 3.5×10^3^ s for the classical Rust Bloom filter. The gap was even larger at query time, where Super Bloom required about 3.1×10^3^ s versus about 6.4×10^3^ s, 1.6×10^4^ s, and 2.0×10^4^ s, respectively. These gains are consistent with the design of Super Bloom, which amortizes block selection across consecutive *k*-mers that share a minimizer.

**Figure 6.**
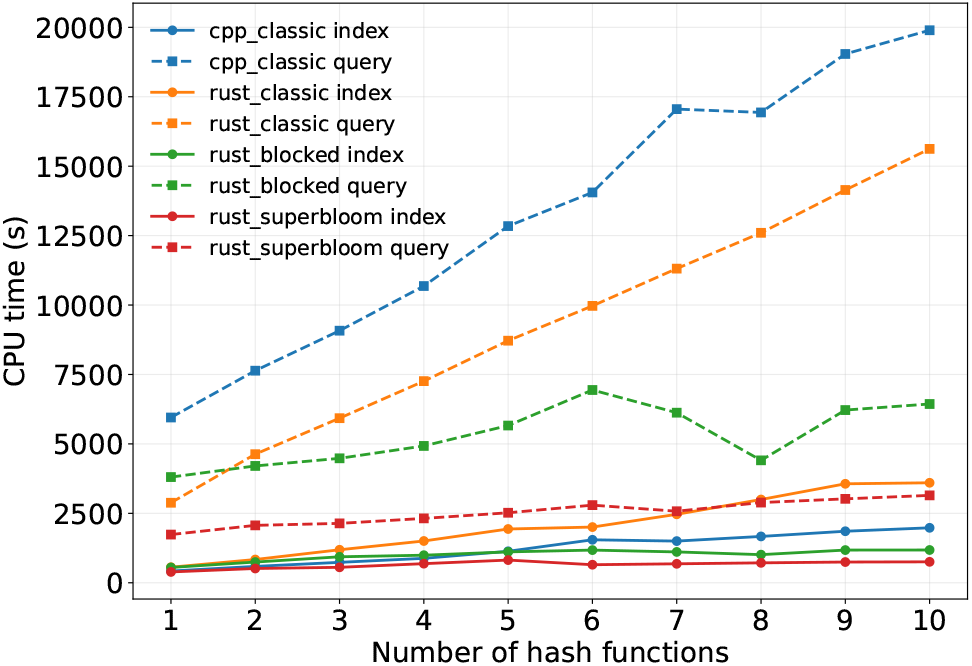
Query CPU time of BioBloom Tools implementations as a function of the number of hash functions on human data, using an index constructed from the human reference genome and 10× simulated human 10 kb reads for querying. The original C++ implementation is compared with a Rust rewrite using either a classical Bloom filter, a blocked Bloom filter, or our novel Super Bloom filter.

The false-positive benchmark on one billion random *k*-mers shows the expected accuracy trade-off (see Figure 7). The C++ and Rust classical Bloom filters are nearly indistinguishable, confirming that the rewrite preserves the behavior of the original structure. The blocked Bloom filter has the highest false-positive count at a fixed memory budget, while Super Bloom without findere (*s* = *k* = 31) remains substantially better than the blocked variant. Adding findere changed this picture markedly. Reducing *s* from 31 to 30, 28, or 24 shifts the curves downward by several orders of magnitude, especially in the low-memory regime. In particular, for *s* = 30 and 2^30^ bits, we observe no false positives among 10^9^ random query *k*-mers. This should be interpreted as an empirical upper bound at the scale of the benchmark rather than as proof of a zero false-positive rate.

**Figure 7.**
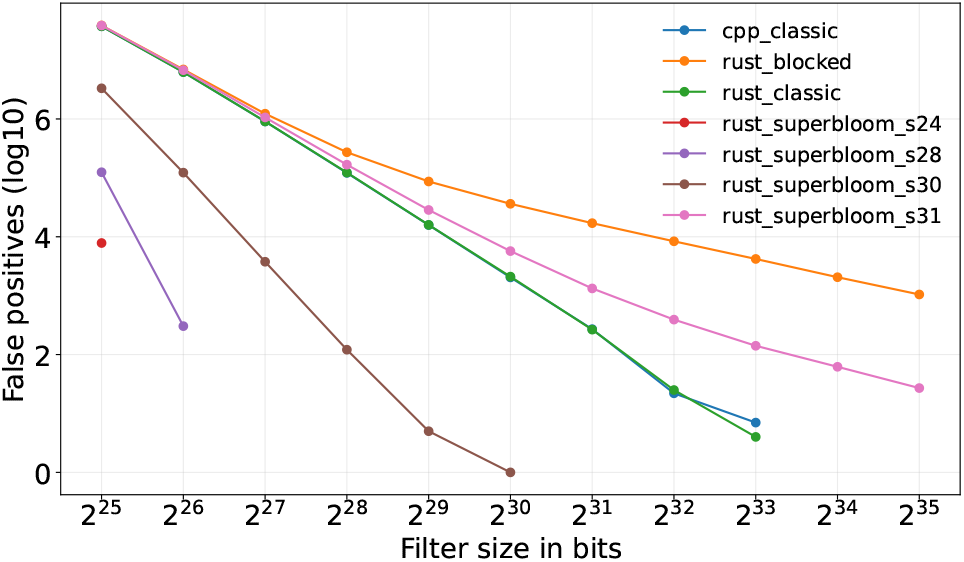
Number of false positives among 10^9^ random *k*-mers as a function of filter size for the different BioBloom Tools implementations on human data, using an index constructed from the human reference genome. The original C++ implementation is compared with a Rust rewrite using either a classical Bloom filter, a blocked Bloom filter, or our novel Super Bloom filter.

### 3.2. State-of-the-art filters

We next compare Super Bloom with several existing Bloom filter implementations: [fastbloom](https://crates.io/crates/fastbloom), generic-bloom, [bloomfx](https://crates.io/crates/bloomfx), bloom-filters, bloom_rs, bloom-filter-rs and roaring-bloom-filter. Roaring Bloom Filter was tested as it is one of the most downloaded Rust implementations of Bloom Filters, but was removed from the results for clarity, as it was an order of magnitude slower than any other tool. Every tool was tested over a range of memory budgets and numbers of hash functions. For insertions, we used the *C. elegans* reference genome WBcel235, and we tested queries by creating reads of 10kb with a 0.1% error rate at 20x coverage.

We first varied the allocated Bloom filter size while keeping the other parameters fixed (see Figure 8 and Figure 9). Across the full range from 2^25^ to 2^38^ bits, Super Bloom remains the fastest implementation both in elapsed time and in total CPU time. Its elapsed time stays particularly low, increasing only from about 1 s to about 7 s, whereas the competing implementations range from about 5–25 s at the smallest sizes to about 28–85 s at the largest sizes. The same ranking is observed in total CPU time, where Super Bloom rises only from about 35 s to about 200 s, while the alternatives span roughly 40–230 s at low memory and up to about 370–750 s at the largest tested sizes. This weak dependence on the memory budget is consistent with the intended locality properties of the structure: enlarging the filter does not substantially degrade the cost of streaming queries when consecutive *k*-mers continue to be processed within the same local block.

**Figure 8.**
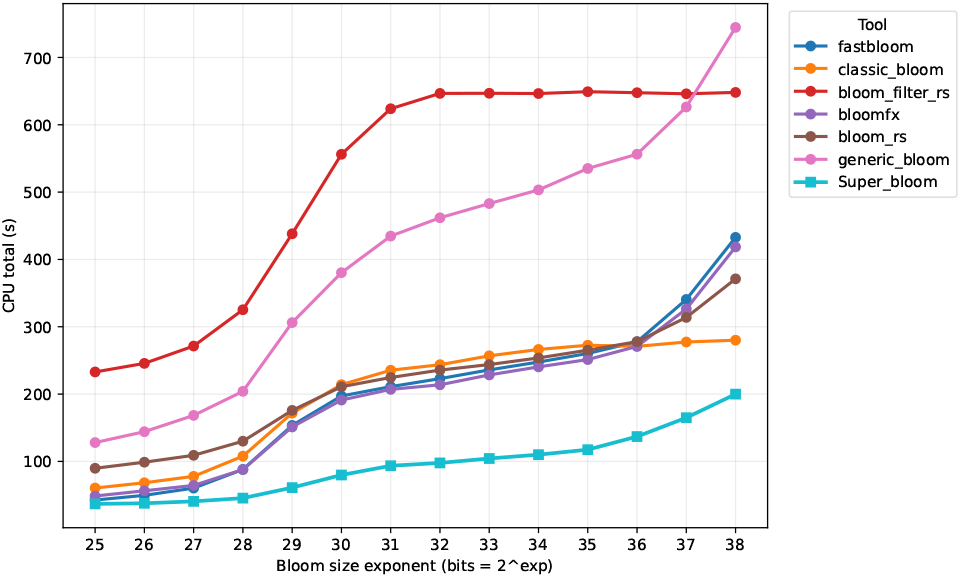
Query CPU time of the different filter implementations as a function of allocated memory on *C. elegans* data, using an index constructed from the reference genome and 10× simulated 10 kb reads for querying.

**Figure 9.**
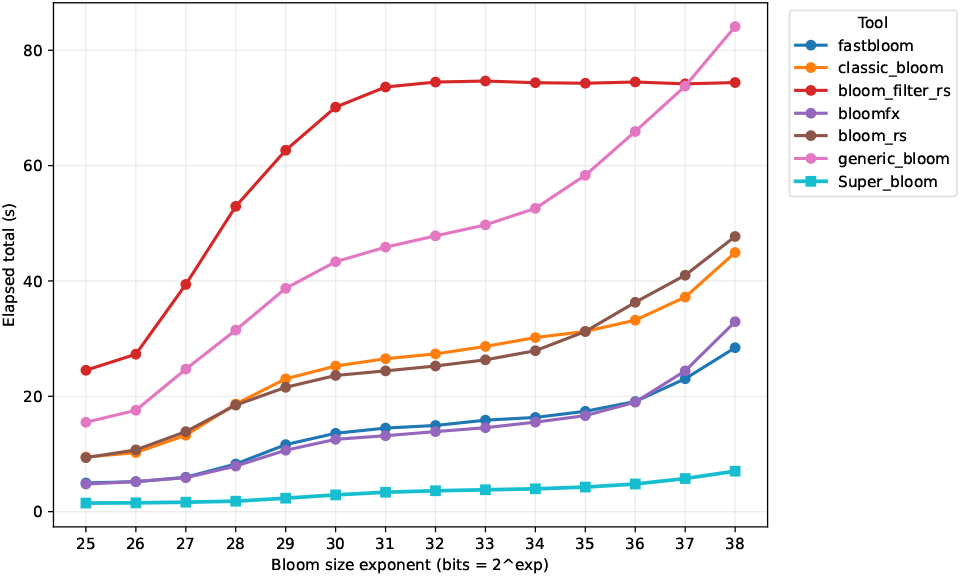
Query elapsed time of the different filter implementations as a function of allocated memory on *C. elegans* data, using an index constructed from the reference genome and 10× simulated 10 kb reads for querying.

We then varied the number of hash functions from 1 to 10 (see Figure 10 and Figure 11). As expected, all implementations become slower when more hashes are used, but the increase is markedly smaller for Super Bloom. Its elapsed time remains nearly flat, growing only from about 6 s to about 8 s, whereas the other implementations rise from about 20–45 s to about 50–190 s. The same trend appears for total CPU time: Super Bloom increases only from about 160 s to about 240 s, while the best competitors reach about 400–560 s and the slowest exceed 800–1800 s. This shallow slope indicates that the cost of additional hash probes is partly amortized by the block-local organization induced by minimizer-based grouping.

**Figure 10.**
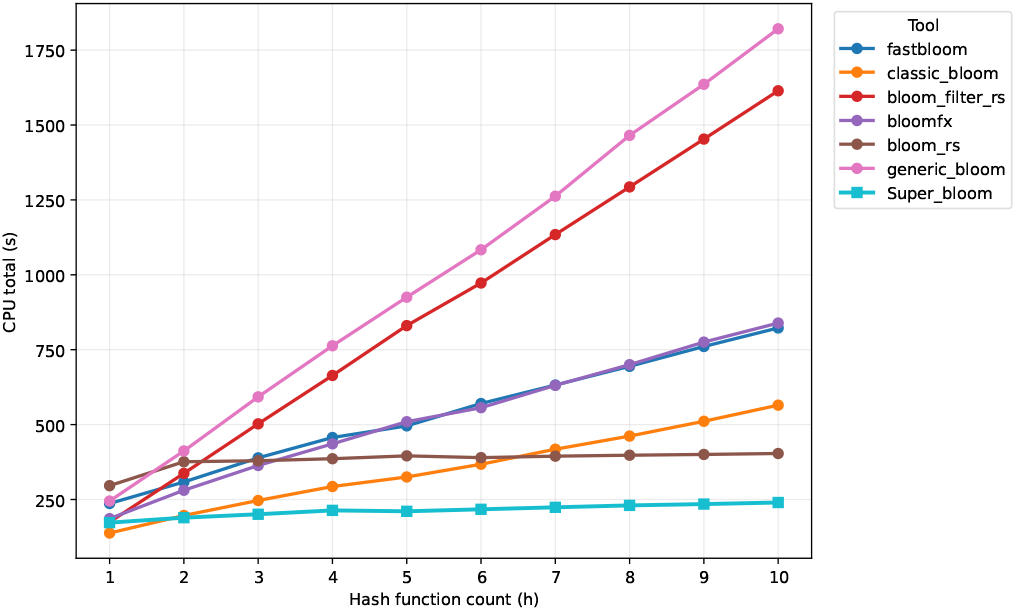
Query CPU time of the different filter implementations as a function of the number of hash functions on *C. elegans* data, using an index constructed from the reference genome and 10× simulated 10 kb reads for querying.

**Figure 11.**
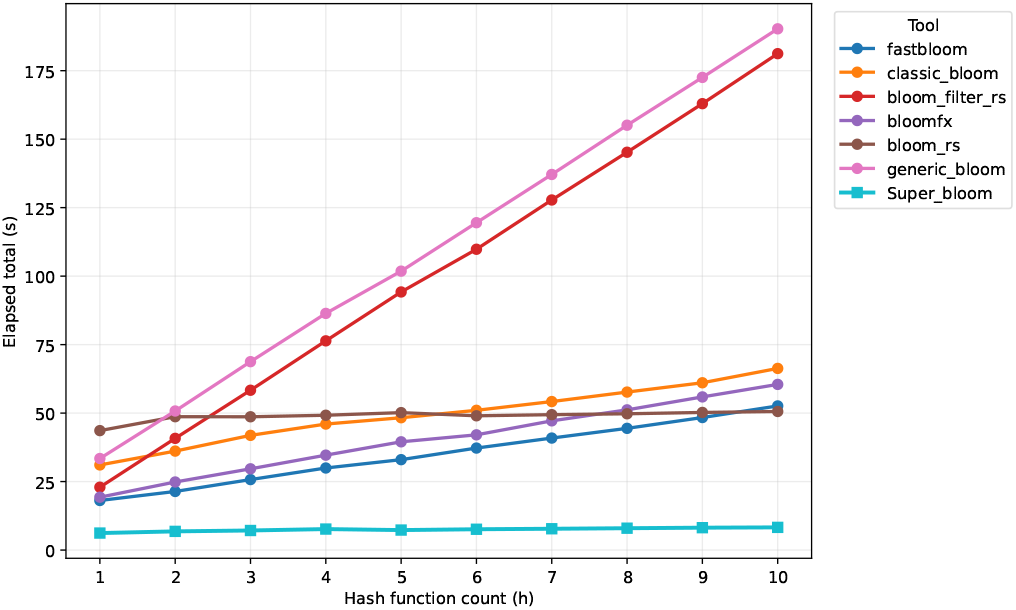
Query elapsed time of the different filter implementations as a function of the number of hash functions on *C. elegans* data, using an index constructed from the reference genome and 10× simulated 10 kb reads for querying.

We also examined the effect of block size at fixed memory (see Figure 12). The wall-clock time depends on the chosen block size, showing that block granularity remains an important tuning parameter even when the total memory budget is held constant. The comparison between a fixed *h* = 8 and a block-size-dependent optimal value of *h* further indicates that the best performance is obtained when the number of hash functions is tuned jointly with the block size rather than chosen independently. Additionally, this figure shows that, for a fixed total number of insertions, runtime begins to increase once the block size reaches 2^{12}^ bits, as indicated by the orange curve no longer remaining flat. This is consistent with the fact that blocks of that size no longer fit within a single cache line, so each access requires loading several cache lines instead of just one.

**Figure 12.**
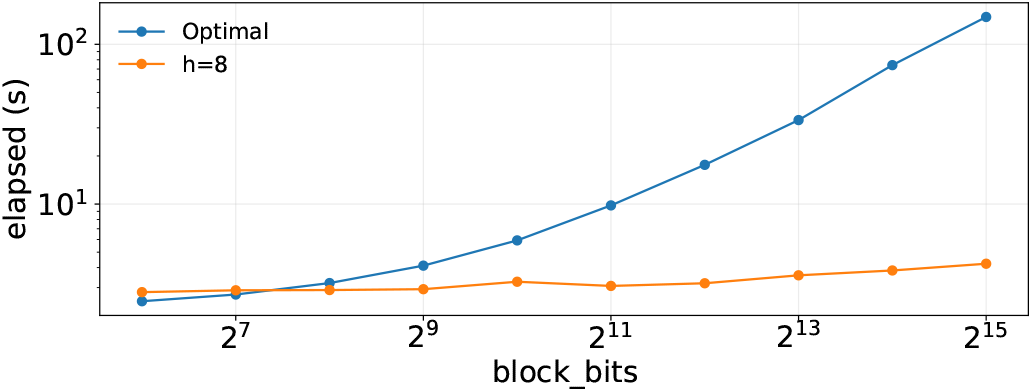
Query wall-clock time on *C. elegans* data as a function of block size at fixed memory (16 GiB). For the “Optimal” curve, the number of hash functions *h* is chosen separately for each block size to maximize block occupancy, whereas for the “*h* = 8” curve it is fixed at 8 across all block sizes.

We also evaluated false positives on 10^9^ random query *k*-mers for a fixed *k* = 31 while varying both the memory budget and the findere parameter *s* (see Figure 13). Without findere, Super Bloom already improves over several standard Bloom-filter implementations at intermediate and large memory budgets, but the main effect comes from the *s*-mer filtration. Decreasing *s* from 31 to 30, 29, 28, 27, 26, or 25 shifts the false-positive curves downward by several orders of magnitude. In particular, the configurations with *s* = 29 or *s* = 30 reach the experimental floor at large memory budgets, while the classical implementations still retain measurable false positives. These results confirm that Super Bloom recovers the speed benefits of blocked layouts while the findere extension compensates for their usual loss in specificity and can even drive the observed false-positive count to zero at the scale of the benchmark.

**Figure 13.**
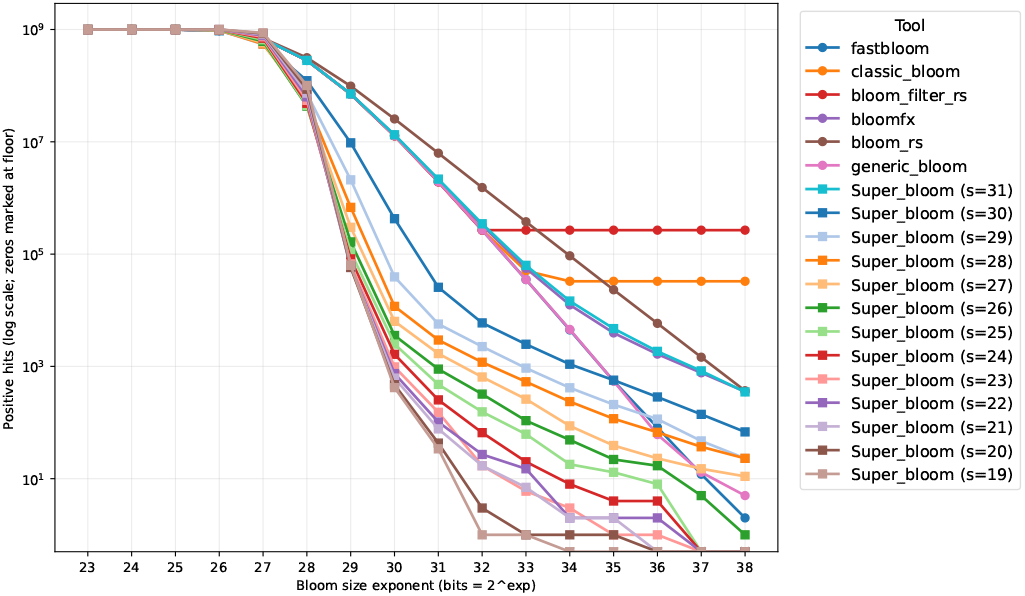
False-positive benchmark on *C. elegans* with 10^9^ random query *k*-mers as a function of filter size. The findere-based Super Bloom variants with *s* < 31 reduce the number of observed false positives by several orders of magnitude, with some settings reaching the experimental floor.

Finally, we measured thread scalability by separating the index and query phases (see Figure 14). For indexing, Super Bloom is already the fastest implementation on a single thread and remains so throughout the full range up to 32 threads, with elapsed time decreasing from about 12 s to about 2 s. For queries, the gain is even stronger: elapsed time drops from about 5 s on one thread to well below 1 s at high thread counts, while the other implementations remain consistently slower. Most tools exhibit diminishing returns beyond a small number of threads, whereas Super Bloom continues to scale well at higher thread counts. This behavior is consistent with a design in which most of the work is local, streaming-friendly, and therefore well suited to parallel execution.

**Figure 14.**
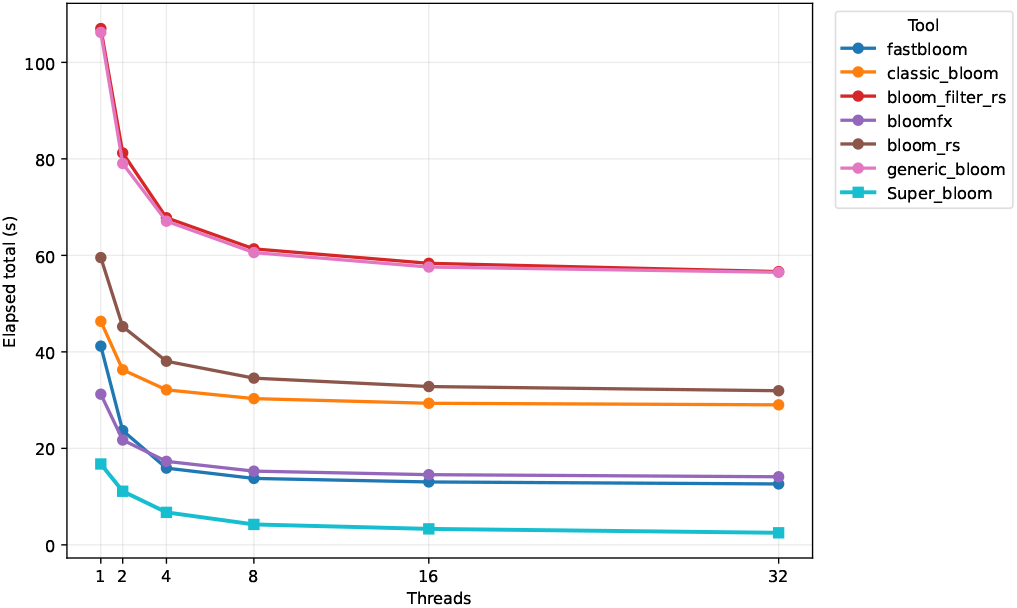
Thread-scaling benchmark on *C. elegans* data, using the reference genome for indexing and 10× simulated 10 kb reads for querying, with 16 GiB of memory and *h* = 4. Left: indexing phase. Right: query phase. Super Bloom is the fastest implementation across the tested thread counts and scales well in both phases.

## 4. Conclusion

In this work, we introduced Super Bloom, a blocked Bloom-filter variant tailored to streaming *k*-mer queries on biological sequences. By mapping all *k*-mers of a super-*k*-mer to the same memory block, Super Bloom converts sequence overlap into improved memory locality and lower random-access cost during sequence-wise processing. Combined with block-level *s*-mer filtration inspired by findere, it achieves both faster queries and substantially lower false-positive rates. Our analysis provides practical parameterization rules, and our experiments show that these gains translate to both stand-alone benchmarks and a BioBloom Tools-like screening workflow.

Several improvements could further strengthen the method. First, the current design relies on standard minimizer schemes, but alternative schemes with lower density could increase average super-*k*-mer length and improve locality even further. Second, the block layout suggests room for more compact representations. When the memory budget is large relative to the practical block size, many blocks remain empty so explicit encodings for empty blocks or compressed representations for sparse ones may reduce memory usage without changing the query model. Third, overloaded blocks remain a critical case, since they weaken the intended false-positive guarantees. One possible extension would be to detect such blocks during construction and associate them with a supplementary local structure, in a way reminiscent of scalable Bloom filters, so that heavily loaded regions could retain a controlled false-positive rate. However, this would also weaken one of the main advantages of the present design, namely its fixed and highly local memory-access pattern, since exceptional blocks would require additional indirections or heterogeneous storage. A key challenge would therefore be to design such a fallback mechanism while preserving cache efficiency and keeping the common-case query path as regular and local as possible.

Beyond these direct optimizations, several research directions appear promising. One natural extension is to adapt such a technique to other filter data structures, such as the counting Bloom filter, in order to support multiplicities, deletions, or adaptive enrichment. Similar questions also arise for quotient-filter-based structures and their improved variants, where locality already plays a central role. Studying whether super-*k*-mer grouping can further improve update cost, cache efficiency, multi-threaded behavior, or practical false-positive control in such designs would be a natural continuation of this work. Another is the static setting, where many bioinformatics workflows naturally separate construction from querying. In that regime, it would be interesting to investigate whether minimizer-based partitioning can be combined with static filters or locality-preserving exact dictionaries, in the spirit of LPHash, to obtain structures that are both more compact and more cache-efficient. Other tools with which to consider these techniques are inverted and bit-sliced indexes, where the dominant cost comes from slice reads and intersections rather than per-filter random accesses. Determining whether minimizer-level grouping can reduce slice bandwidth, improve compressibility, or enable earlier pruning in these settings remains an open question.

Another promising option to consider is the extension of the framework beyond contiguous *k*-mers to more mutation-tolerant seeds, including spaced seeds[103], [104], [105], gapped *k*-mers[106], and strobemers[107]. Such seeds are attractive because they can improve robustness to substitutions or more complex mutation patterns, but they do not come with an obvious analogue of the minimizer-based grouping used here. In particular, HackGap[106] explicitly notes that the minimizer-based bucketing strategy of fast contiguous *k*-mer methods does not transfer easily to gapped *k*-mers, while strobemers derive their sensitivity from a more composite seed structure. A natural question is therefore whether one can define, for these seed families, a locality-preserving gathering rule that would play the role of minimizers for contiguous *k*-mers and still support the same cache-efficient block organization. Finding such a rule remains an open problem.

More broadly, Super Bloom suggests that overlap-aware filter design is a promising path to explore for sequence bioinformatics, and that approximate membership structures can benefit substantially from exploiting the non-independence of consecutive *k*-mers rather than treating them as isolated keys.

## 7. Acknowledgments

The authors are grateful to Igor Martayan for helpful technical discussions.

## Appendix A

**Block occupancy distribution by number of super-*k*-mers**

In this section, we compare the observed block-occupancy distribution with the Poisson approximation derived in the main text (see Section 2.5). More precisely, Table 1, Table 2, Table 3, and Table 4 report the numbers of blocks containing respectively 0, 1, 2, and at least 3 super-*k*-mers, for multiple total numbers of blocks. Overall, the agreement is good across the tested range, with the largest deviations appearing in intermediate regimes.

**Table 1.** Observed and predicted numbers of empty blocks as a function of the total number of blocks when indexing the *C. elegans* reference genome WBcel235. Predictions are obtained from the Poisson approximation derived in the main text.

| Total blocks | Observed | Poisson prediction | Relative gap | Observed proportion (%) |
| --- | --- | --- | --- | --- |
| 32768 | 0 | 0 | 0 % | 0 % |
| 65536 | 0 | 0 | 0 % | 0 % |
| 131072 | 0 | 0 | 0 % | 0 % |
| 262144 | 0 | 0 | 0 % | 0 % |
| 524288 | 0 | 0 | 0 % | 0 % |
| 1048576 | 0 | 0 | 0 % | 0 % |
| 2097152 | 1344 | 720 | 86.67 % | 0.06 % |
| 4194304 | 109419 | 77754 | 40.72 % | 2.61 % |
| 8388608 | 1343531 | 1131848 | 18.7 % | 16.02 % |
| 16777216 | 6702052 | 6153365 | 8.92 % | 39.95 % |
| 33554432 | 21178556 | 20290368 | 4.38 % | 63.12 % |
| 67108864 | 53235531 | 52106697 | 2.17 % | 79.33 % |
| 134217728 | 119362921 | 118089247 | 1.08 % | 88.93 % |
| 268435456 | 252763115 | 251410699 | 0.54 % | 94.16 % |
| 536870912 | 520176199 | 518782662 | 0.27 % | 96.89 % |
| 1073741824 | 1055319303 | 1053904601 | 0.13 % | 98.28 % |

**Table 2.**
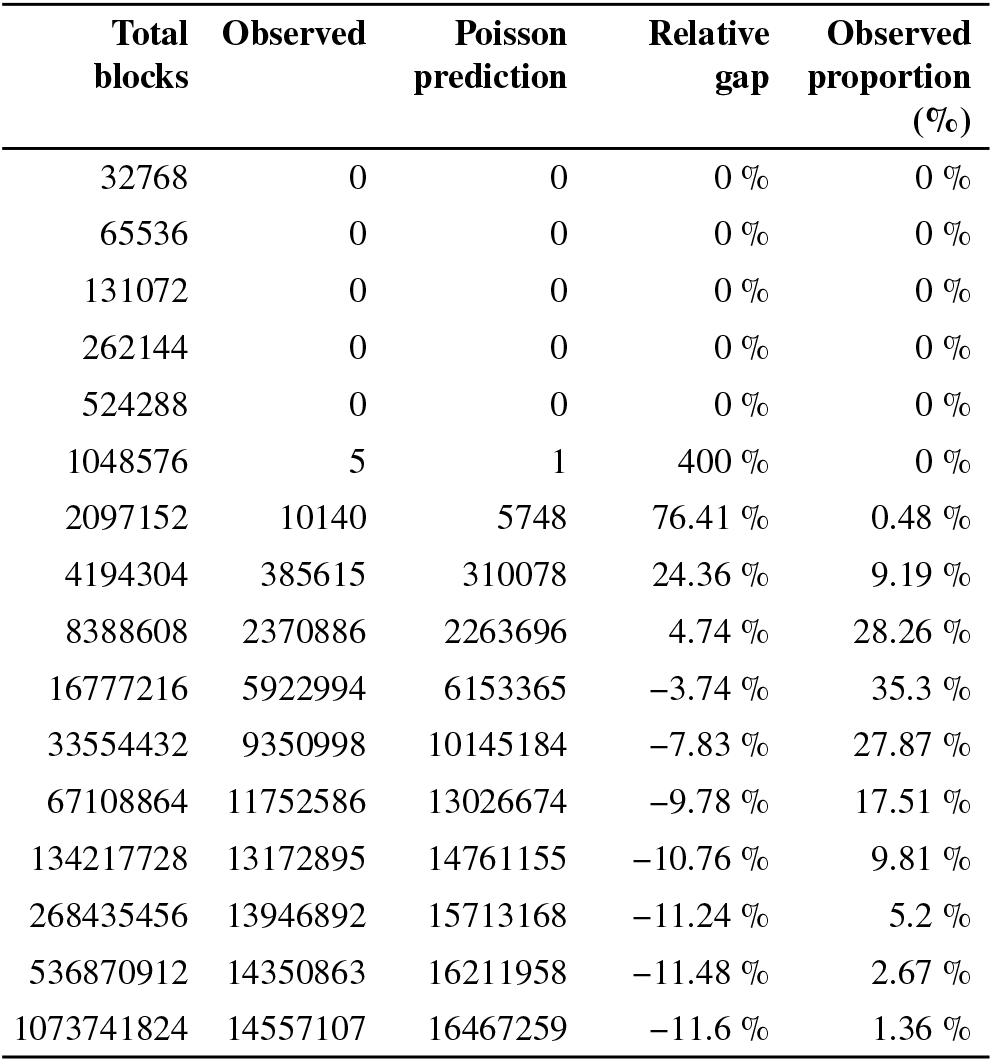
Observed and predicted numbers of blocks containing exactly one super-*k*-mer as a function of the total number of blocks when indexing the *C. elegans* reference genome WBcel235. Predictions are obtained from the Poisson approximation derived in the main text.

**Table 3.** Observed and predicted numbers of blocks containing exactly two super-*k*-mers as a function of the total number of blocks when indexing the *C. elegans* reference genome WBcel235. Predictions are obtained from the Poisson approximation derived in the main text.

| Total blocks | Observed | Poisson prediction | Relative gap | Observed proportion (%) |
| --- | --- | --- | --- | --- |
| 32768 | 0 | 0 | 0 % | 0 % |
| 65536 | 0 | 0 | 0 % | 0 % |
| 131072 | 0 | 0 | 0 % | 0 % |
| 262144 | 0 | 0 | 0 % | 0 % |
| 524288 | 0 | 0 | 0 % | 0 % |
| 1048576 | 49 | 15 | 226.67 % | 0 % |
| 2097152 | 35597 | 22923 | 55.29 % | 1.7 % |
| 4194304 | 688026 | 618286 | 11.28 % | 16.4 % |
| 8388608 | 2157464 | 2263696 | -4.69 % | 25.72 % |
| 16777216 | 2760898 | 3076682 | -10.26 % | 16.46 % |
| 33554432 | 2296957 | 2536296 | -9.44 % | 6.85 % |
| 67108864 | 1587490 | 1628334 | -2.51 % | 2.37 % |
| 134217728 | 1053824 | 922572 | 14.23 % | 0.79 % |
| 268435456 | 731350 | 491036 | 48.94 % | 0.27 % |
| 536870912 | 554783 | 253311 | 119.01 % | 0.1 % |
| 1073741824 | 462272 | 128650 | 259.33 % | 0.04 % |

**Table 4.**
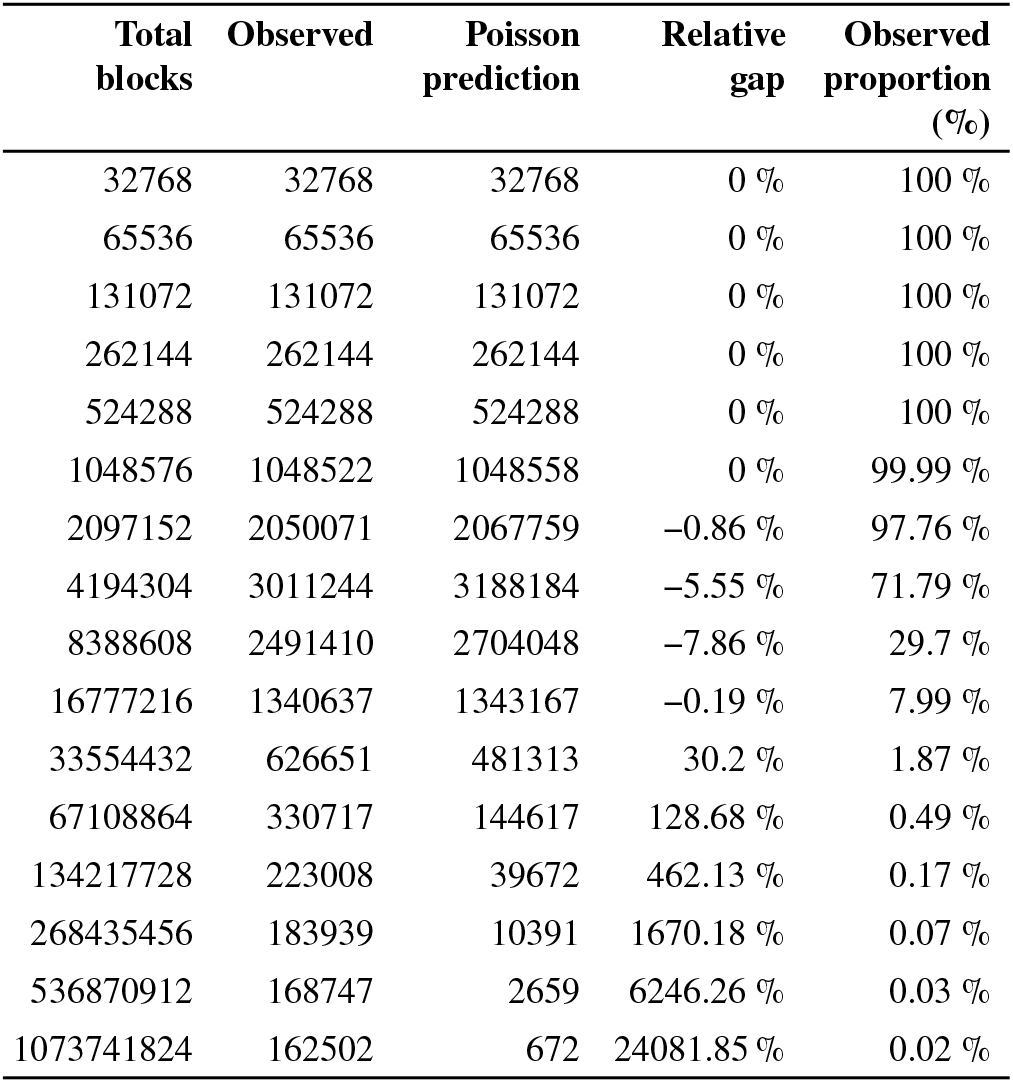
Observed and predicted numbers of blocks containing at least three super-*k*-mers as a function of the total number of blocks when indexing the *C. elegans* reference genome WBcel235. Predictions are obtained from the Poisson approximation derived in the main text.

| Total blocks | Observed | Poisson prediction | Relative gap | Observed proportion (%) |
| --- | --- | --- | --- | --- |
| 32768 | 32768 | 32768 | 0 % | 100 % |
| 65536 | 65536 | 65536 | 0 % | 100 % |
| 131072 | 131072 | 131072 | 0 % | 100 % |
| 262144 | 262144 | 262144 | 0 % | 100 % |
| 524288 | 524288 | 524288 | 0 % | 100 % |
| 1048576 | 1048522 | 1048558 | 0 % | 99.99 % |
| 2097152 | 2050071 | 2067759 | -0.86 % | 97.76 % |
| 4194304 | 3011244 | 3188184 | -5.55 % | 71.79 % |
| 8388608 | 2491410 | 2704048 | -7.86 % | 29.7 % |
| 16777216 | 1340637 | 1343167 | -0.19 % | 7.99 % |
| 33554432 | 626651 | 481313 | 30.2 % | 1.87 % |
| 67108864 | 330717 | 144617 | 128.68 % | 0.49 % |
| 134217728 | 223008 | 39672 | 462.13 % | 0.17 % |
| 268435456 | 183939 | 10391 | 1670.18 % | 0.07 % |
| 536870912 | 168747 | 2659 | 6246.26 % | 0.03 % |
| 1073741824 | 162502 | 672 | 24081.85 % | 0.02 % |

## Appendix B

**Effect of the findere scheme on false positives for random negative and one-mismatch-neighbor *k*-mers**

In this section, we evaluate how the findere parameter *s* affects the empirical false-positive rate for two classes of queries: random negative *k*-mers and one-mismatch neighbors of indexed *k*-mers.

As discussed in Section 2.6, a one-mismatch neighbor differs from an indexed *k*-mer only at its last base. When queried with the findere scheme, its first *z* = *k* − *s* overlapping *s*-mers are shared with the indexed *k*-mer and are therefore truly present in the filter. Consequently, in this one-mismatch-last-base setting, only the final queried *s*-mer must be falsely reported present for the whole *k*-mer to be reported present. By contrast, a random negative *k*-mer typically requires positive answers for all *z* + 1 = *k* − *s* + 1 constituent *s*-mers.

To isolate the effect of *s*, we indexed the *E. coli* reference genome ASM584v2 for several values of *s* and chose *h* so that (*z* + 1) · *h* = 12 in every configuration. This keeps the nominal number of tested hash positions per queried *k*-mer constant across configurations.

Figure 15 shows the expected trade-off. As *z* increases, the falsepositive rate of one-mismatch neighbors increases because fewer false-positive events are needed on the non-shared part of the query. In contrast, the false-positive rate of random negative *k*-mers remains nearly constant, with a slight decrease that is consistent with a lower effective fill ratio when overlapping *k*-mers share more inserted *s*

**Figure 15.**
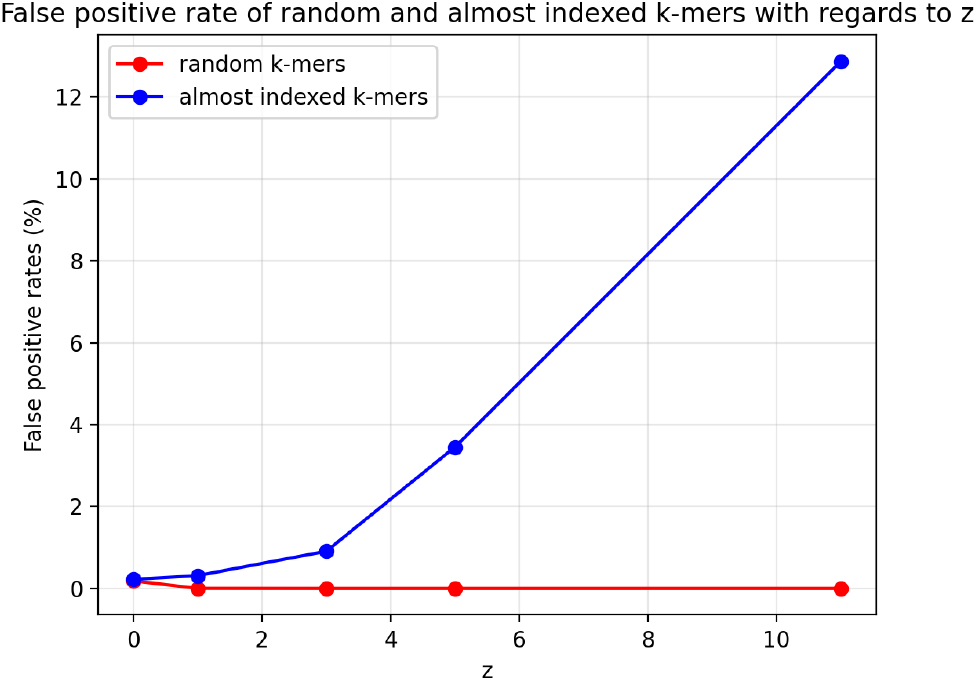
Empirical false-positive rates of random negative and one-mismatch-neighbor *k*-mers as a function of *z* = *k* − *s*, after indexing the *E. coli* reference genome ASM584v2 in a filter of size 2^26^ bits. For each value of *z, h* was chosen so that (*z* + 1) · *h* = 12.

